# MELATONIN INDUCES CHANGES ON THE CIRCADIAN RHYTHMS OF REPRODUCTIVE HORMONES DURING SPERMATOGENESIS IN PUBERTAL MALE SEA BASS, *Dicentrarchus labrax*

**DOI:** 10.1101/2024.07.16.603757

**Authors:** M.Victoria Alvarado, Felipe Espigares, Manuel Carrillo, Alicia Felip

## Abstract

Reproduction is a highly demanding biological process that occurs at the optimal time of the year and day to ensure the success of spawn and offspring. Melatonin is a hormone that, secreted mainly by the pineal gland, plays a critical role in the integration of the photoneuroendocrine information from environment (annual and daily variations) to modulate reproductive activity and gonadal development in fish. In this study we assessed the effect of exogenous melatonin on the circadian levels of sex steroids and gonadotropins in pubertal 2 yr-old male sea bass during their reproductive cycle including, pre-spermatogenesis (Pspg), spermiation (Spm) and post-spermiation (PSpm) stages. Our results demonstrated that all reproductive hormones displayed circadian variations along the entire reproductive cycle in pubertal fish. Circulating levels of the luteinizing hormone (Lh) were affected by both melatonin injection and the daily timing of administration during the Spm and PSpm stages, thus evoking variations of Lh levels at night. Melatonin also significantly affected circadian rhythms of Fsh during the Spm stage. Overall, both 11-Kt and T plasma levels displayed circadian variations during the reproductive cycle in the sea bass which were not prevented by melatonin. However, melatonin showed a significant decrease of plasma levels of 11-Kt 1h after dusk during the Pspg stage, while it increased those levels of T 5 h after dusk during the PSpm stage. These findings provide new insights into the role of melatonin in fish reproduction as a key factor in regulation of daily variation of key hormones involved in gonadal development. This circumstance may have implications in the control of gametogenesis and management of fish in aquaculture.

**Highlights:**

- Plasma levels of Fsh, Lh, T and 11-Kt show daily rhythms in pubertal male sea bass.
- Melatonin evokes changes in daily rhythms of Lh during the reproductive cycle.
- Melatonin yields Fsh plasma level differences during spermiation stage.
- Melatonin elicits T and 11-Kt plasma level differences depending on reproductive stage and time.

## 1. Introduction

Environmental factors, such as light and temperature, play a crucial role in guiding animals to adopt the most suitable life strategy to optimize their reproductive success and ensure the survival of offspring. Melatonin, the major hormone responsible for integrating both annual and circadian cues in the organism, is synthesized by the pineal gland and eyes and acts as an endogenous clock regulating some biological processes such as reproduction in vertebrates (Li et al., 2023; Makris et al., 2023) including fish species (Falcón et al., 2010, 2011; Lombardo et al., 2014; Maitra and Hasan, 2016; Guellard et al., 2019; Krylov et al., 2021). It is known that brain melatonin receptor activation regulates gonadotropin-releasing hormone (GnRH) gene expression and secretion, and consequently the secretion of pituitary gonadotropins as well as the production of steroids in the gonads in fish (Hofman, 2006; Parhar et al., 2012; Múñoz-Cueto et al., 2020). Many studies have focused on the melatonin system in fish due to its involvement in the control of circadian variations of key endocrine factors and the regulation of reproduction (Migaud et al., 2010; Carrillo et al., 2015; Maitra and Hasan, 2016; Cowan et al., 2017). Recent evidence shows that melatonin has considerable potential for optimizing animal physiological functions and, in turn, improving the efficiency of animal husbandry (Li et al 2023; Contreras-Correa et al., 2023).

Melatonin, that is closely associated with the day night cycle, has been extensively investigated in a large number of fish since it has been observed that it is involved in the photoneuroendocrine regulation of reproduction in species such as European sea bass (*Dicentrarchus labrax*) (Bayarri et al., 2004, 2010), Mozambique tilapia (*Oreochromis mossambicus*) (Nikaido et al., 2009) and the broadhead catfish (*Clarias macrocephalus*) (Aripin et al., 2014) among others. Furthermore, evaluation of the effects of exogenous administration of melatonin -by implants, injection or bath treatments- on reproductive system in fish has showed that this hormone is involved in ovarian follicle maturation and fecundity in zebrafish (*Danio rerio*) (Carnevali et al., 2011) and modules both GnRH and kisspeptin systems in European sea bass (Servili et al., 2013; Alvarado et al., 2015). In addition, the presence of melatonin in fish seminal plasma suggests a melatonin role in fish reproductive antioxidant system, thus preventing and counteracting oxidative stress damage on fish spermatozoa (Félix et al., 2020). These findings indicate that variations in circulating melatonin levels might affect the daily rhythms of reproductive hormones during gametogenesis. Indeed, environmental cycles, which exhibit both seasonal and day/night changes of light and temperature with an annual and daily periodicity, contribute to synchronize reproductive rhythms in fish that display daily spawning and egg viability rhythms. Thus, spawning events occur near or during the night in nocturnal species, whereas it takes place during the day in diurnal species (Villamizar et al., 2012). These insights open up new opportunities for a better understanding of the endocrine mechanisms regulating fish reproduction and, in this way improve those protocols and practices in laboratory conditions and fish hatcheries (Blanco-Vives and Sánchez-Vázquez, 2009; Villamizar et al., 2012).

The European sea bass is a highly valued marine fish species which is considered a short-day breeder. In the Mediterranean region, sea bass reproduces under short photoperiod regimes during the winter (December-March) and ceases in the spring (April), when the day length increases (Carrillo et al., 2009; Felip and Piferrer, 2018). This species exhibits nocturnal locomotor activity rhythms during the reproductive season, although it is diurnal during the resting period, with spawning events occurring at night (Villamizar et al., 2012). In addition, sea bass exhibits a daily profile of demand-feeding activity that changes seasonally (Sánchez-Vázquez et al., 1998), thus showing a predominantly diurnal feeding pattern with the greatest activity being concentrated at the end of the day (Azzaydi et al., 1999). Males usually mature at 2-yr of age, although early onset puberty affects 20-30% of 1 yr-old males in the population under intensive culture conditions (Felip and Piferrer, 2018). The use de continuous light (LL) from early stages of development reduces the incidence of precocious males (Begtashi et al., 2004; Felip et al., 2008; Rodríguez et al., 2012). Under LL, daily variations in plasma melatonin are disrupted in males, although melatonin binding variations persist in hypothalamus during the entire reproductive cycle, from pre-spermatogenesis to post-spermiation, including both spermatogenesis and spermiation stages (Bayarri et al., 2010). Interestingly, the effect of photoperiod on daily rhythms of melatonin, reproductive hormones and gonadal development on caged sea bass has demonstrated that artificial lights suppress the circulating plasma levels of melatonin and significantly affect the daily rhythm of luteinizing hormone (Lh) storage and release (Bayarri et al., 2004). The aim of this study was to analyse the effects of melatonin on the daily rhythms of sex steroid and gonadotropins in pubertal males of 2 yr-old during critical stages of their reproductive cycle including pre-spermatogenesis, spermiation and post-spermiation stages. The crosstalk existing between melatonin and these main reproductive hormones proves that changes of circadian rhythms are critical in sea bass reproduction and, thus the use of chronobiology across aquaculture systems might make fish production more efficient.

## 2. Material and Methods

### 2.1. Animal housing

A total of a hundred and twenty 2-yr old adult European male sea bass (average body weight of 254.43 ± 54.52 g) were obtained from Cupimar (Cádiz, Spain) and maintained under natural photoperiod and seawater temperature conditions (12-23 ± 1°C) at the Institute of Aquaculture Torre de la Sal (IATS, Castellón, Spain, 40°N 0°E) facilities. Fish were distributed in six 500 L circular fiberglass tanks (n = 20 fish/tank). Animals were fed once a day until apparent satiety with a commercial dry pellet diet supplied by Proaqua Nutrición, S.A. (Palencia, Spain) (protein 54–45%, lipids 20–12%, carbohydrates 9–25%, ash 11%, moisture 1–3%, DE 22.4-19.7 MJ kg−1).

### 2.2. Experimental design and sampling

Fish were organized into two groups for each of the three critical gametogenesis stages considered in this study based on gonad growth and development of this species: i) pre-spermatogenesis (PSpg), referred to as spermatogonial proliferation and renewal (September); ii) spermiation (Spm), associated with testicular maturation and sperm production (January) and iii) post-spermiation (PSpm), referred to as resting (April) (Bayarri et al., 2010). Two tanks were established in each gametogenesis stage with a first group, acted as the control (Control, C), in which the animals were injected with a saline solution (NaCl 0.65%) and a second group in which the animals were intraperitoneally injected with melatonin dissolved in vehicle solution (saline solution containing 1% ethanol) (Melatonin, M). A single dose of 0.5 μg melatonin (Sigma, St. Louis, MO) per gram of fish body weight was used. Animals were injected at dusk coinciding with the sunset in each stage including PSpg at 19:30 h, Spm at 17:55 h and PSpm at 20:50 h. The dose used in this study was based on previous research conducted in this species (Servili et al., 2013). Fish were anesthetized with MS-222 (0.1 g l^-1^ of sea water) (Sigma-Aldrich; Merck group, Darmstadt, DE) before each sampling and injection procedure. All animals were handled according to the guidelines for animal experiments established by the Spanish Royal Decree (RD 53/2013) and European legislation (Directive 2010/63/EU) for the use of laboratory animals for scientific purposes.

### 2.3. Hormonal analysis

Blood samples were collected from the caudal vein at four different times in each stage during the night: 1 hour after dusk coinciding with the time at which melatonin was injected, 5 hours after dusk, 2 hours before dawn, and at the sunrise. Plasma was separated by centrifugation at 4 °C and stored at − 20 °C until analysed. Plasma sex steroids were measured by conventional enzyme immunoassay (EIA) which were validated for sea bass in our laboratory. Circulating testosterone (T) plasma levels were measured according to the method described by Rodríguez et al. (2000a). The sensitivity of the assay was 9.0 pg/ml (Bi/B0 = 94%), with inter- and intra-assay coefficients of variability of 9.5% and 6.2%, respectively. Levels of 11-ketotestosterone (11-Kt) were measured according to Rodríguez et al. (2001). The sensitivity of the assay was 7.0 pg/ml (Bi/B0 = 90%), with inter- and intra-assay coefficients of variability of 9.5% and 10%, respectively. Levels of sea bass luteinizing hormone (Lh) (Mateos et al., 2006) and follicle-stimulating hormone (Fsh) (Molés et al., 2012) were measured by a homologous competitive enzyme-linked immunosorbent assays (ELISA) developed for this species. The assay sensitivity of Lh was 0.65 pg/ml (Bi/B0 = 80%) with inter- and intra-assay coefficients of variability of 11% and 11.7%, respectively. The assay sensitivity of Fsh was 500 pg/ml (Bi/B0 = 93.9%) with inter- and intra-assay coefficients of variability of 5.4% and 2.12%, respectively.

### 2.4 Statistical analysis

Data are presented as the mean ± the standard error of the mean (SEM). Plasma hormone levels for each gametogenesis stage were analyzed using a two-way ANOVA (SigmaStat 3.5), followed by the Holm-Sidak test for post hoc multiple comparisons. Data were transformed to meet normality and homoscedasticity requirements, as needed. Plasma levels of 11-Kt and T were log-transformed in both PSpg and Spm stages, whereas Fsh and Lh levels were square-root transformed in the PSpg and in both Spm and PSpm stages, respectively. Statistical significance was set at P < 0.05 (Sokal and Rohlf, 1981). Outliers were identified using the generalized extreme studentized deviate procedure, with a significance level of P = 0.05 and a maximum of 20% outliers per sample size (GraphPad Prism 10.0).

## 3. Results

### 3.1 Effects of melatonin on circulating plasma levels of reproductive hormones during pre-spermatogenesis (PSpg)

Fsh levels were significantly affected by melatonin administration (p = 0.016) and the time after treatment injection (p < 0.001) during the PSpg stage, although the interaction between variables was not significant (P = 0.137) (Table 1). These findings indicated that circulating plasma levels of Fsh showed a daily variation during the PSpg stage, although melatonin did not alter its circadian rhythm (Fig. 1A). By contrast, circulating plasma Lh levels showed statistically significant differences in the interaction between the melatonin administration and the time post-injection (P = 0.037) (Table 1). In this sense, fish in the control group exhibited a progressive but not significant increase of Lh from 1 hour after dusk to 2 hours before down, with a maximum Lh concentration occurring at 05:30 h. Then, plasma Lh levels significantly decreased at dawn (Fig. 1B). Of note, fish in the melatonin group showed a progressive but not significant increase of Lh levels from dusk to dawn, when circulating Lh levels were significantly higher than those of the controls (Fig. 1B). These results demonstrated that melatonin evoked changes in the daily plasma Lh levels in this species during PSpg stage. Regarding circulating plasma levels of androgens, the control group exhibited significant daily variations of 11-Kt, with significant high levels recorded at 20.30 h (1h after melatonin injection) although dropped drastically 5 hours after dusk and remained unchanged until 2 hours before dawn (Fig. 1C). 11-Kt concentration in melatonin-injected fish was significantly lower than those of control fish 1 hour after dusk and the daily variation of this hormone was significantly affected by the time post-injection and its interaction with treatment administrated during PSpg (Table 1). On the other hand, variation of the circulating plasma levels of T were not affected by melatonin. The time post-injection elicited significant changes in the T during the PSpg stage, although the interaction with the treatment received by fish was similar between groups (Fig. 1D) (Table 1).

**Figure 1.**
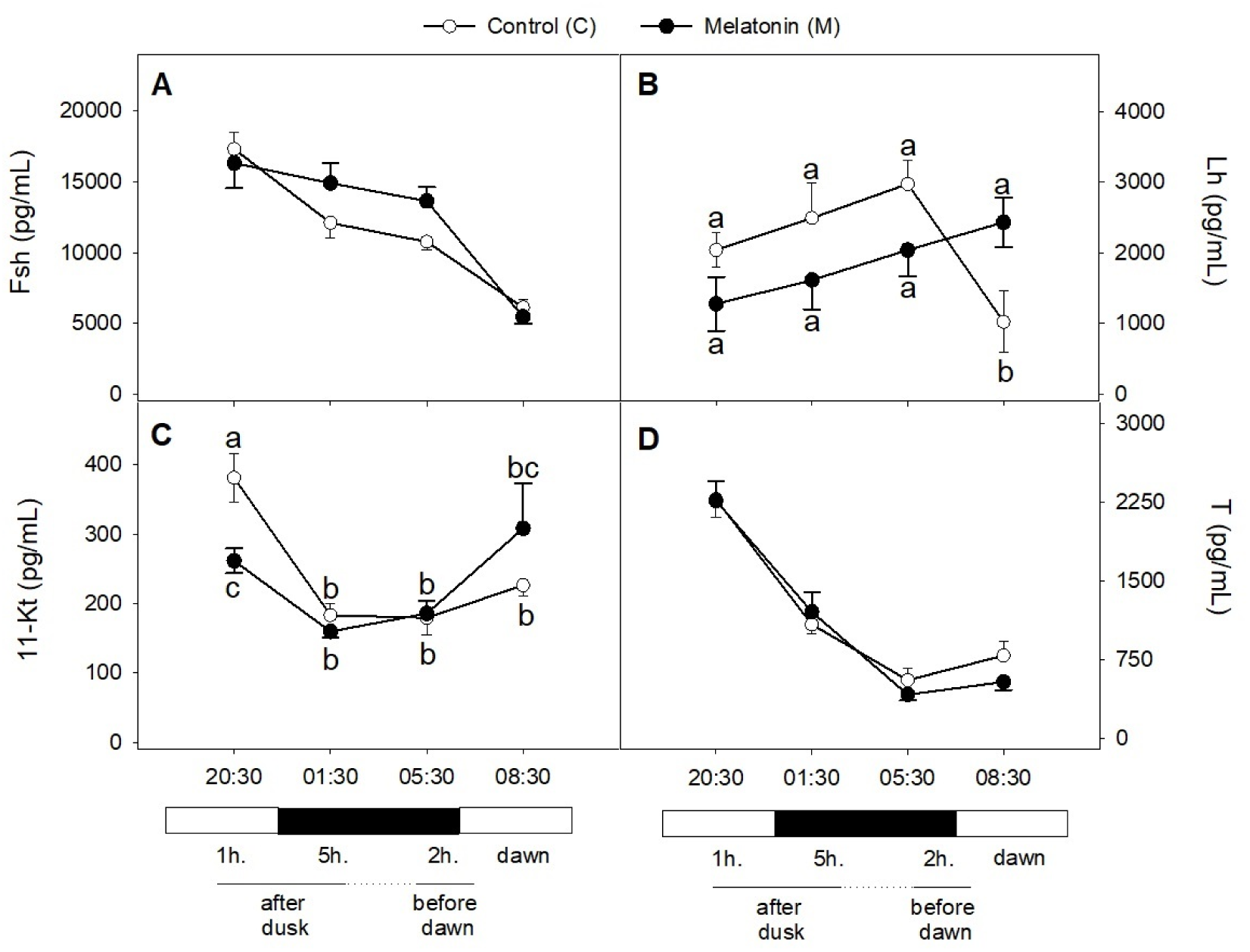
Daily plasma levels of follicle-stimulating hormone (Fsh) (A), luteinizing hormone (Lh) (B), 11-Ketotestosterone (11-Kt) (C) and Testosterone (T) (D) of 2-yr old male European seabass during pre-spermatogenesis (PSpg). Control group (white points) was injected saline and experimental group (black points) was injected melatonin. Data are expressed as the mean ± SEM. Letters indicate significant differences (P < 0.05) between both groups in the same sample point and over the time in each group (n= 7-21 fish/group).

**Table 1.** Results of the two-way ANOVA tests to assess the effect of treatment (control vs melatonin), time post-injection (1 hour after dust vs 5 hours after dust vs 2 hours before dawn vs dawn) and the interaction between both variables on plasma levels of gonadotropins (Fsh, Lh) and sex steroids (11-Kt, T) during pre-spermatogenesis (Pspg), spermiation (Spm) and post-spermation (PSpm) stage. Significant differences (P < 0.05) are indicated in bold.

| Hormone | Main effects and interactions | Pspg |  | Spm |  | PSpm |  |
| --- | --- | --- | --- | --- | --- | --- | --- |
|  |  | F-value | P-value | F-value | P-value | F-value | P-value |
| Fsh | Treatment | 6.087 | **0.016** | 4.360 | **0.041** | 7.027 | **0.010** |
|  | Time post-injection | 17.706 | **< 0.001** | 7.300 | **< 0.001** | 1.351 | 0.265 |
|  | Treatment x Time post-injection | 1.909 | 0.137 | 4.561 | **0.006** | 0.300 | 0.825 |
| Lh | Treatment | 1.073 | 0.304 | 6.465 | **0.013** | 33.915 | **< 0.001** |
|  | Time post-injection | 2.116 | 0.106 | 16.445 | **< 0.001** | 13.965 | **< 0.001** |
|  | Treatment x Time post-injection | 2.989 | **0.037** | 8.046 | **< 0.001** | 24.055 | **< 0.001** |
| 11-Kt | Treatment | 0.222 | 0.639 | 0.0825 | 0.775 | 3.527 | 0.065 |
|  | Time post-injection | 19.384 | **< 0.001** | 16.770 | **< 0.001** | 4.563 | **0.006** |
|  | Treatment x Time post-injection | 2.904 | **0.041** | 0.474 | 0.702 | 0.315 | 0.815 |
| T | Treatment | 0.922 | 0.340 | 0.865 | 0.356 | 12.288 | **< 0.001** |
|  | Time post-injection | 47.106 | **< 0.001** | 17.698 | **< 0.001** | 49.629 | **< 0.001** |
|  | Treatment x Time post-injection | 0.476 | 0.700 | 1.654 | 0.185 | 9.920 | **< 0.001** |

### 3.2 Effects of melatonin on circulating plasma levels of reproductive hormones during spermiation (Spm)

Circulating plasma levels of Fsh (Fig. 2A) and Lh (Fig. 2B) showed daily variations during the Spm stage that were altered by melatonin. The daily variations in circulating levels of both gonadotropins showed that Fsh and Lh levels were significantly affected by melatonin and the time post-injection (Table 1). In addition, plasma levels of Fsh (P = 0.006) and Lh (P < 0.001) showed a statistically significant interaction between these variables, thus evoking that melatonin-injected fish attained the highest values of Fsh 5 hours after dusk (Fig. 2A), whereas Lh decreased 5 hours after dusk and remained unchanged until 2 hours before dawn (Fig. 2B). On the other hand, daily variation in circulating levels of T (Fig. 2B) and 11-Kt (Fig. 2C) were similar to those of the melatonin-injected fish during the Spm stage, although significant differences were recorded affecting the time post-injection (Table 1).

**Figure 2.**
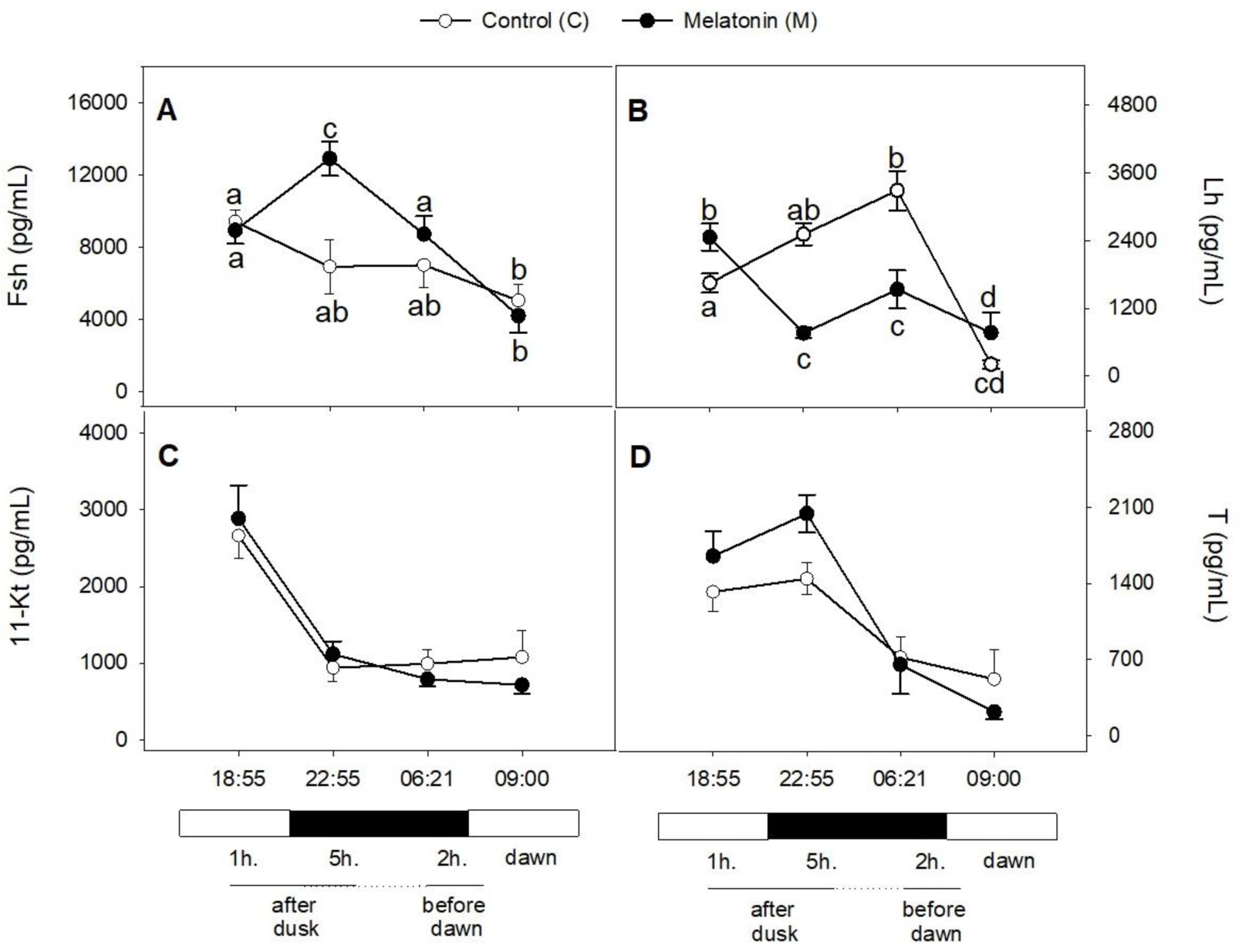
Daily plasma levels of follicle-stimulating hormone (Fsh) (A), luteinizing hormone (Lh) (B), 11-Ketotestosterone (11-Kt) (C) and Testosterone (T) (D) of 2-yr old male European seabass during spermiation (Spm). Groups and representation of data as described in Figure 1.

### 3.3 Effects of melatonin on circulating plasma levels of reproductive hormones during post-spermatogenesis (PSpm)

Fsh levels were significantly affected by melatonin administration during PSpm stage, although the interaction between both treatment and timepost-injection variables did not show statistical differences (Fig. 3A) (Table 3). On the other hand, a daily variation was observed in the circulating levels of Lh that showed a statistically significant interaction between these two variables. Fish in the control group displayed a decrease in plasma Lh levels 5 hours after dusk which were maintained unchanged during dusk followed by a significant increase at dawn (Fig. 3B). Melatonin-injected animals showed unchanged levels from 1 to 5 hours after dusk, although a significant increase was observed at 05:05 h (2 hours before down) and, then decreased at dawn. In addition, daily variations of 11-Kt (Fig. 3C) were significantly affected by the time after melatonin administration (Table 1), whereas plasma levels of T in the control group displayed a slight increase from 1 hour after dusk to 2 hours before dawn, followed by a smooth decrease at dawn (Fig. 3D). Significant daily variation in plasma T was found in melatonin-injected fish, with a peak value being recorded at 01:50 h during dusk, followed by a progressive decrease from 5 hours after dusk to dawn. Levels of T were significantly affected (P < 0.001) by melatonin administration, the time post-injection and its interaction (Table 1).

**Figure 3.**
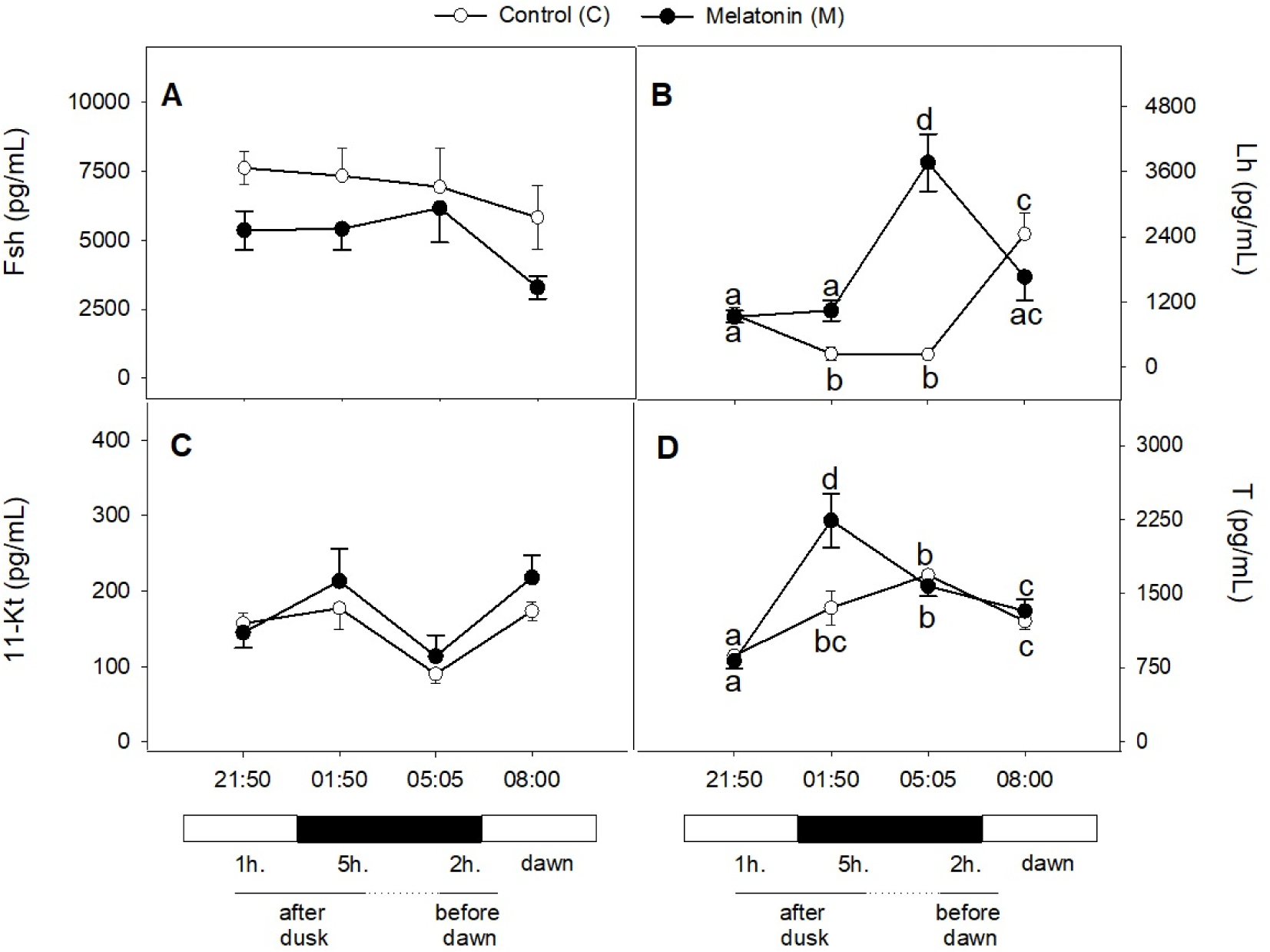
Daily plasma levels of follicle-stimulating hormone (Fsh) (A), luteinizing hormone (Lh) (B), 11-Ketotestosterone (11-Kt) (C) and Testosterone (T) (D) of 2-yr old male European seabass during post-spermiation (PSpm). Groups and representation of data as described in Figure 1.

## 4. Discussion

This study provides physiological insights into the actions of melatonin on the daily variations of circulating levels of gonadotrophin hormones and sexual steroids of male sea bass at the onset of puberty throughout their entire seasonal reproductive cycle. Our results agree with previous studies on sea bass providing evidence for the role that melatonin has in the endocrine control of reproduction of this species. In these sense, the long-term melatonin exposure, administrated as an implant, elicits seasonal changes in key reproductive hormones of males that affect testicular maturity in adult fish (Alvarado et al., 2015). On the other hand, intraperitoneally injected melatonin has been documented to evoke inhibitory effects in the brain, thus affecting the expression of *gnrh-1*, *gnrh-3* and several *gnrh* receptors in this teleost fish (Servili et al., 2013). Furthermore, the absence of the melatonin rhythm in juvenile fish, caused by continuous light regimes, effectively reduces the incidence of male precocity (Begtashi et al., 2004; Bayarri et al., 2004, 2009). Of note, this study reports, for the first time, circadian variation of Fsh throughout the entire reproductive cycle of 2 yr-old male sea bass and that it is affected by melatonin during Spm stage.

Our results demonstrate that endogenous melatonin secretion negatively affects circulating plasma Fsh levels, which show a nocturnal tendency to decrease, thus displaying an opposite diurnal rhythm compared to that of the Lh that exhibits an increasing nocturnal variation. Interestingly, exogenous melatonin raises nocturnal Fsh levels until the sunrise in contrast to the decrease in Lh observed during Spm, thus supporting the idea that melatonin has a role in the endocrine control of spermatogenesis in the sea bass. In this sense, it is known that plasma Lh exhibits a circadian variation in juvenile male sea bass with a significant increase of nocturnal Lh plasma concentration during critical stages of spermatogenesis under natural photoperiod conditions. In fact, the lack of plasma Lh daily rhythmicity under continuous light, at some points of gonadal development, has been considered a possible reason to explain the inhibition of precocity under this photoperiod regime in this species (Bayarri et al., 2009). In fact, it has been documented that the hormonal cascade is prevented from being completed in juvenile fish reared under continuous light photoperiod (Rodríguez et al., 2019). It is worthy to note that those disturbances affecting plasma Lh variations in juveniles have been also observed in males during their second year of age in this study. Pubertal male sea bass displayed a nocturnal rise of Lh during the dark phase that then dropped at the sunrise during PSpg and Spm, whereas daily rhythm of Lh dropped during darkness and increase at sunrise during PSpm. These seasonal changes in Lh plasma levels are considered to be crucial at the onset of puberty in this teleost fish, as the slight increases of this hormone coincide with important reproductive stages -PSpg and Spm- in which the proliferation of spermatogonia occurs (Rodríguez et al., 2005; Espigares et al., 2015). Accordingly, the nocturnal melatonin levels are considered to induce a positive regulation of the Lh levels at nighttime in the sea bass to promote the progression of gametogenesis (Bayarri et al., 2004; Falcon et al 2010; Carrillo et al., 2015). Of note, the high concentration of plasma Lh levels in controls during the dark phase throughout these periods of time support the already suggested involvement of Lh in the regulation of gamete maturation and spermiation in fish (Swanson et al., 2003). Interestingly, the administration of exogenous melatonin evokes changes of these circadian rhythms of Lh levels, a fact that suggests that this hormone influences the testicular growth and development in the sea bass. Accordingly, differences in circadian variations of both gonadotropins might be related with the fact that differential regulation mechanisms exist for Fsh and Lh release throughout Gnrh in fish (Karigo et al., 2013). Regarding diurnal Fsh variations, changes have been also observed in male humans and rams suggesting that these circadian rhythms might be crucial for the maintenance of the physiological functions of the testis (Brambilla et al., 2009; Zhang et al., 2018; Samir et al., 2022). Effects of melatonin on testicular function and spermatogenesis in bulls have evidenced for an important role of this hormone to improve the efficiency of livestock reproduction (Li et al., 2023). Recent studies in teleosts have revealed that melatonin acts as a hormone and an antioxidant in the control of fish reproduction (Félix et al., 2020; Takahashi and Ogiwara, 2021). Melatonin is reported to exert its antioxidant activity to reduce the free radical damage in the ovary and improve the quality of oocytes (Maitra and Hassan, 2016), whereas it may contribute to antioxidant capacity of seminal plasma or positively impact on different fish sperm quality biomarkers including sperm concentration, motility and velocity (Félix et al., 2023ab). Therefore, further research should be conducted on the role of melatonin as a potential agent for regulation of spermatogenesis and preserve sperm quality in vertebrates (Ebrahimi et al, 2023; Makris et al., 2023).

Regarding circadian variations of sex-steroids, the detection of maximum concentrations of 11-Kt were observed during Spm stage as previously described in this species (Bayarri et al., 2009), thus reinforcing the role that 11-Kt plays in the progress of spermatogenesis in fish (Rodriguez et al., 2000b; Rodriguez et al., 2005; Higuchi et al., 2017). Although melatonin does not prevent circadian variation of plasma 11-Kt, it is worthy to note that the lowest values of this androgen are recorded when fish are kept under continuous light, in other words when the melatonin rhythm is disturbed as a consequence of this photoperiod regime (Bayarri et al., 2009). Similarly to 11-Kt, T shows a circadian variation during the reproductive cycle in the sea bass, which is not prevented by melatonin. Collectively, these findings support the role that melatonin has on the reproductive axis of sea bass at the brain level as previously described in this species (Servili et al., 2013; Alvarado et al., 2015). It is known the exogenous melatonin modulates transcriptional actions of the daily variations of gnrh system in the brain of sea bass (Servili et al., 2013), although its putative involvement in the circadian modulation of kisspeptin system are still to be investigated. Nevertheless, evidence exists that long-term melatonin administration induces the downregulation of kisspeptin-gnrh members in the brain that affects *fhsβ* transcription and, in turn, disturbing testicular maturity in the sea bass (Alvarado et al., 2015). Accordingly, further studies are necessary in order to decipher those mechanisms that mediate physiological actions of melatonin in the brain and gonads of teleost fish species, as it may contribute to make fish production more efficient.

## 5. Conclusion

In summary, our study reveals that the main reproductive hormones display circadian variation in male sea bass throughout the entire seasonal reproductive cycle during their second year of life. Of note, exogenous melatonin seems to evoke physiological actions in the pituitary gland, thus affecting differentially gonadotropin secretion throughout gametogenesis. Melatonin yielded changes of circadian variation of plasma Lh during Pspg, Spm and PSpm, whereas Fsh plasma concentration differences were observed during Spm stage. On the other hand, melatonin did not prevent circadian variation of plasma T and 11-KT (Fig.4).

**Figure 4.**
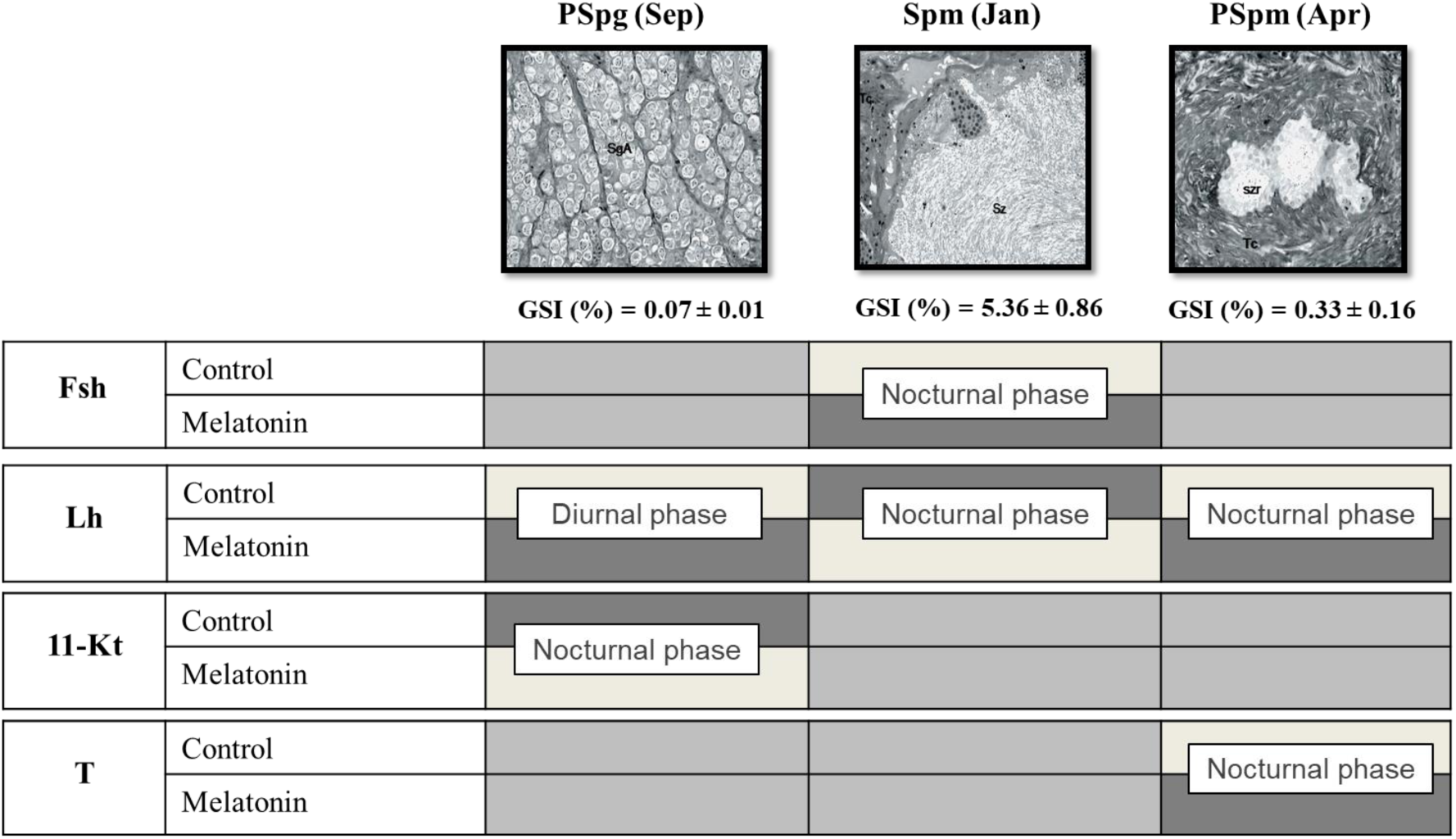
Summary of physiological changes of the circadian rhythms of reproductive hormones in 2 yr-old male sea bass subjected to exogenous melatonin (by injection) during their entire reproductive season including three critical gametogenesis stages: i) pre-spermatogenesis (PSpg), referred to as spermatogonial proliferation and renewal (September); ii) spermiation (Spm), associated with testicular maturation and sperm production (January) and iii) post-spermiation (PSpm), referred to as resting (April). The shading in each box represents the levels of the variable being considered: light grey, significantly low levels; grey-intermediate, levels with no significant differences and dark grey, significantly high levels.

## ACKNOWLEDGMENTS

We thank to Histological Services and Animal Husbandry at IATS. This work was financially supported by grants KISSCONTROL (AGL2009-11086) from the Spanish Ministry of Science and Innovation (MICINN) and the Spanish Ministry of Economy and Competitiveness (MINECO), and REPROBASS (PROMETEO/2010/003) and CIAICO/2022/002 from the Valencian Regional Government. M.V. Alvarado was supported by FPI fellowship (AGL2010-036443) from the MICINN and MINECO.

## AUTHOR CONTRIBUTIONS

M.V.A.: conceptualization, data curation, formal analysis, methodology, investigation, validation, visualization, writing—original draft and writing—review and editing; F.E.: data curation, formal analysis, investigation, validation and writing—review and editing; M.C.: conceptualization, formal analysis, funding acquisition, project administration, resources, supervision, visualization, writing—original draft and writing—review and editing; A.F.: conceptualization, funding acquisition, project administration, resources, supervision, visualization, writing—original draft and writing—review and editing. All authors gave final approval for publication and agreed to be held accountable for the work performed therein.

## DATA AVAILABILITY

Data is made available as supplementary material to the manuscript.

